# An autoregulation loop in *fust-1* for circular RNA regulation in *Caenorhabditis elegans*

**DOI:** 10.1101/2021.03.22.436400

**Authors:** Dong Cao

## Abstract

Circular RNAs (circRNAs) are always expressed tissue-specifically, suggestive of specific factors that regulate their biogenesis. Here, taking advantage of available mutation strains of RNA binding proteins (RBPs) in *Caenorhabditis elegans*, I performed a screening of circRNA regulation in thirteen conserved RBPs. Among them, loss of FUST-1, the homolog of FUS (Fused in Sarcoma), caused downregulation of multiple circRNAs. By rescue experiments, I confirmed FUST-1 as a circRNA regulator. Further, I showed that FUST-1 regulates circRNA formation without affecting the levels of the cognate linear mRNAs. When recognizing circRNA pre-mRNAs, FUST-1 can affect both exon-skipping and circRNA in the same genes. Moreover, I identified an autoregulation loop in *fust-1*, where FUST-1, isoform a promotes the skipping of exon 5 of its own pre-mRNA, which produces FUST-1, isoform b with different N-terminal sequences. FUST-1, isoform a is the functional isoform in circRNA regulation. Although FUST-1, isoform b has the same functional domains as isoform a, it cannot regulate either exon-skipping or circRNA formation.

## Introduction

Although treated as byproducts of splicing in early years (Cocquerelle et al., 1993; Nigro et al., 1991), circRNAs have shown diverse functions in different biological or physiological environments, including interactions with DNAs (transcription regulation (Li et al., 2015), R-loop structure formation (Conn et al., 2017)), RNAs (miRNA sponge (Hansen et al., 2013; Memczak et al., 2013)), and proteins (Du et al., 2020; Okholm et al., 2020; Xia et al., 2018; Zhu et al., 2019). Rather than splicing errors, the circRNA production process is well-regulated, in which both intronic sequences (*cis* elements) and RBPs (*cis/trans* elements) are involved (Chen, 2020). Reverse complementary matches (RCMs, *cis* elements) in introns that flank exon(s) to be circularized promote circRNA formation, presumably by bringing splice sites for back-splicing together. RBPs (*cis*/*trans* elements) can regulate back-splicing positively or negatively. Muscleblind in *Drosophila* promotes the production of the circRNA from the second exon of its own pre-mRNA by binding to the flanking introns (Ashwal-Fluss et al., 2014). The splicing factor Quaking promotes circRNA biogenesis during epithelial to mesenchymal transition (Conn et al., 2015). Immune factors NF90/NF110 promote circRNA formation by associating with intronic RNA pairs in circRNA-flanking introns (Li et al., 2017). RBM20 in mice promotes the production of multiple circRNAs from the gene Tintin (Khan et al., 2016). ADAR1 (Ivanov et al., 2015; Rybak-Wolf et al., 2015) and DHX9 (Aktas et al., 2017) negatively regulate circRNAs by disturbing the base pairing of RCMs. Multiple heterogeneous nuclear ribonucleoproteins (hnRNPs) and serine–arginine (SR) proteins function in a combinatorial manner to regulate circRNAs in cultured *Drosophila* cells (Kramer et al., 2015). HNRNPL regulates circRNA levels in LNCaP cells by binding to circRNA-flanking introns (Fei et al., 2017). A recent paper shows that RBP FUS affects circRNA expression in stem cell-derived motor neurons in mice (Errichelli et al., 2017). All these findings were from *in vitro* cultured cells of different organisms. *C. elegans* provides a suitable animal model for *in vivo* study of circRNA regulation, given the conservation of RBPs and availability of diverse mutant strains.

FUS plays diverse roles in DNA repair and RNA splicing (Sama et al., 2014). Particularly, the mutation of FUS has been linked to the neurodegenerative disease ALS (Kwiatkowski et al., 2009; Vance et al., 2009). *C. elegans* has been used to model ALS by knocking in wild-type or mutated human FUS (Markert et al., 2019; Murakami et al., 2015; Murakami et al., 2012; Vaccaro et al., 2012a; Vaccaro et al., 2012b; Veriepe et al., 2015). As the homolog of FUS in *C. elegans*, FUST-1 is involved in lifespan and neuronal integrity regulation (Therrien et al., 2016) and miRNA-mediated gene silencing (Zhang et al., 2018).

Autoregulation feedback has been found in many RBPs, which is beneficial for them to maintain proper protein levels (Buratti and Baralle, 2011; Muller-McNicoll et al., 2019). The mechanisms include autoregulation of alternative splicing (AS) of their own pre-mRNA, which either produces unproductive transcripts with premature termination codons (PTCs) that are subjected to nonsense-mediated decay (NMD) pathway (McGlincy et al., 2010; Rossbach et al., 2009; Sureau et al., 2001; Wollerton et al., 2004) or produces another protein isoform with disturbed functional domains (Damianov and Black, 2010). Here, I identified an autoregulation pathway in the production of the two isoforms of FUST-1 in *C. elegans*.

Here, using available RBP mutation strains in *C. elegans*, I performed a screening of 13 conserved RBPs in their roles in circRNA regulation. FUST-1 stood out in the screening, showing promotional effects on the production of multiple circRNAs. I further checked FUST-1’s role in circRNA regulation globally by RNA-seq and found FUST-1 can also negatively regulate circRNAs. FUST-1 recognizes pre-mRNAs of circRNA genes and can regulate both exon-skipping and circRNA production in the same genes. Moreover, I characterized an autoregulation loop in the production of the two isoforms of FUST-1, in which FUST-1, isoform a promotes the skipping of exon 5 of *fust-1* pre-mRNA, which produces FUST-1, isoform b. Interestingly, although FUST-1, isoform b has the same functional domains as isoform a, it cannot regulate exon-skipping or circRNA formation.

## Results

### 1. RBP screening identifies FUST-1 as a circRNA regulator

Previous studies have shown that circRNAs are expressed in a tissue-specific and well-regulated manner (Chen and Schuman, 2016; Gruner et al., 2016; Memczak et al., 2013; Rybak-Wolf et al., 2015; Westholm et al., 2014; You et al., 2015), suggesting the existence of specific factors that regulate circRNA production. Here, taking advantage of the available RBP mutants in *C. elegans*, I aimed to identify potential circRNA regulators. Several previously identified circRNAs that were either neuron-enriched or highly expressed in neurons were selected as targets (Figure S1A) (Cao, 2021). Thirteen RBPs that are conserved and have expressions in the neurons were chosen as potential regulators (Norris et al., 2017). Using mutant strains of these RBPs, a screening by RT-qPCR was performed to check the level changes of selected circRNAs in these mutant strains compared with wild-type N2 strain at the L1 stage (Figure 1A). As expected, levels of some circRNAs were altered in these mutant strains. Interestingly, most level changes of the selected circRNAs in these mutants were downregulations, suggestive of these RBPs’ beneficial roles in circRNA production. Moreover, multiple neuron-enriched circRNAs (*circ-glr-2, circ-iglr-3, circ-arl-13, circ-cam-1*) were found to be downregulated in several strains (*asd-1(csb32), tiar-3(csb35), fox-1(csb39), mec-8(csb22), hrpf-1(csb26)*, and *fust-1(csb21)*), suggesting the regulation of these circRNAs by multiple RBPs. This is consistent with their roles in alternative splicing, where combinational regulation of one target by multiple RBPs is common in *C. elegans* (Tan and Fraser, 2017). No additive effect in circRNA regulation was found in *fust-1(csb21); hrpf-1(csb26)* double mutant strain compared with *fust-1(csb21)* single mutation (Figure S1B), suggesting that these RBPs may function as parts of a whole RNA-protein complex.

**Figure 1.**
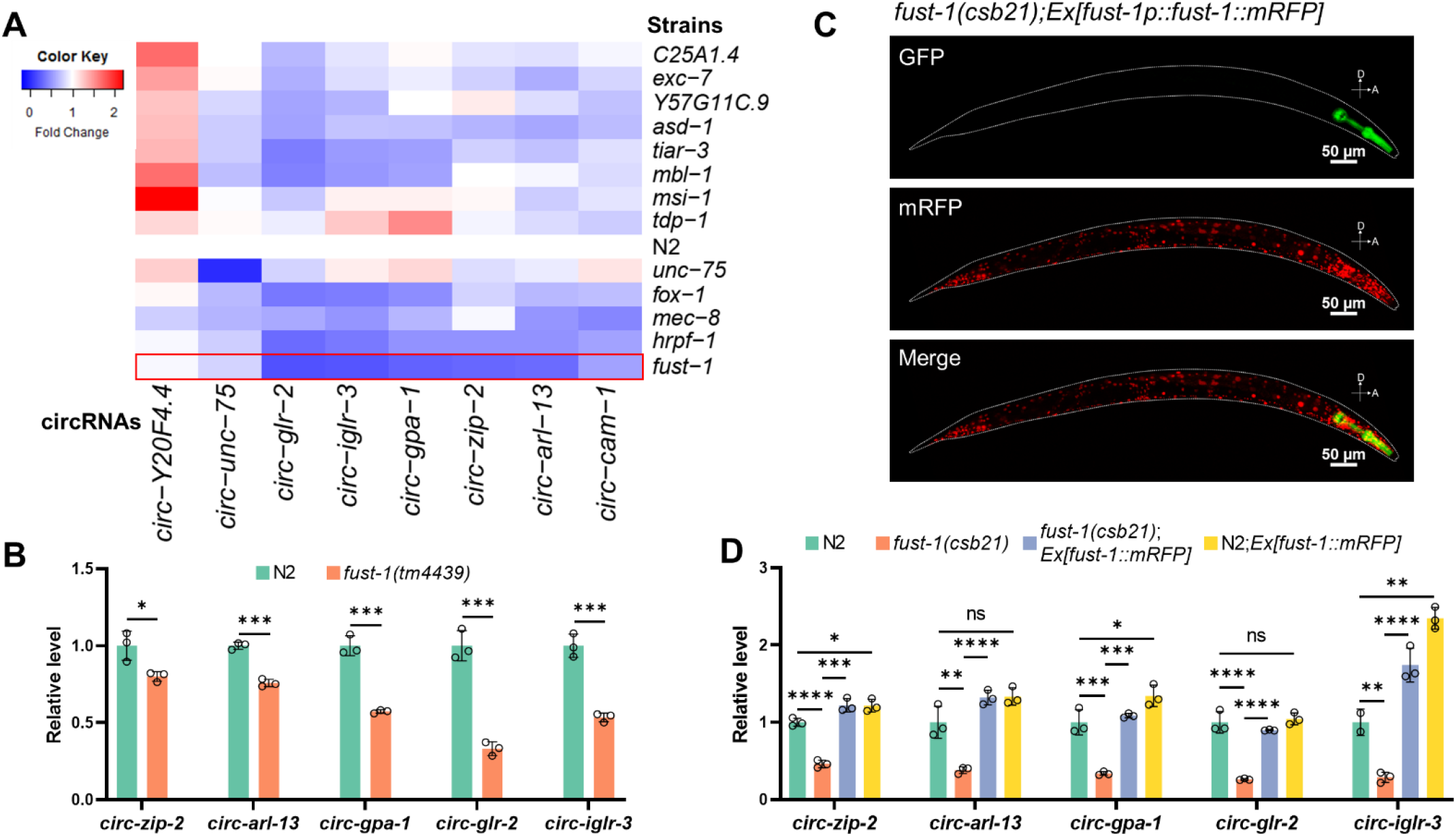
RBP screening identifies FUST-1 as a circRNA regulator. (A) Heatplot showing the fold changes of circRNAs in 13 RBP mutant strains compared with wild-type N2 strain. Foldchanges are quantified by RT-qPCR and normalized to the N2 strain using *pmp-3* as the reference gene. Blue color means downregulation and red color means upregulation. (B) RT-qPCR quantification of circRNA levels in wild-type N2 strain and *fust-1(tm4439)* strain. Levels are normalized to the N2 strain using *pmp-3* as the reference gene. Results are shown as mean ± sd of three biological replicates. Two-tailed Student’s *t*-test. *p*<0.05, \*\**p* < 0.01, \*\*\**p* < 0.001. (C) Representative images showing the expression pattern of mRFP-fused FUST-1 in *fust-1(csb21)* strain. Note the pharyngeal GFP expression in *fust-1(csb21)*. Scale bars: 50 μm. (D) RT-qPCR quantification of circRNAs in the indicated strains. Levels are normalized to the N2 strain using *pmp-3* as the reference gene. Results are shown as mean ± sd of three biological replicates. One-way ANOVA, Tukey’s multiple comparisons. \**p*<0.05, \*\**p* < 0.01, \*\*\**p* < 0.001, \*\*\*\**p* < 0.0001; ns, not significant.

In these strains, *fust-1(csb21)* showed the most substantial downregulation of multiple circRNAs (Figure 1A). Hence it was chosen for further investigation. The downregulation of these circRNAs was also found in another *fust-1* mutant strain *fust-1(tm4439)* (Figure 1B and Figure S1C). To further confirm the role of *fust-1* in circRNA regulation, a rescue strain (*fust-1(csb21); Ex[fust-1::mRFP]*) and an overexpression strain (N2; *Ex[fust-1::mRFP]*) were made with extrachromosomal expression of *fust-1* genomic sequence, starting from *fust-1* promoter (2181 bp upstream ATG) to just before the stop codon of *fust-1*. Monomeric red fluorescent protein (mRFP) was fused to the C-terminal to check expression patterns. The expression of FUST-1 was mainly in the nucleus of neurons and intestinal cells (Figure 1C and S1D). The mRFP-positive L1 worms from the rescue strain and the overexpression strain were sorted, and levels of the circRNAs were checked by RT-qPCR. As expected, the levels of downregulated circRNAs were restored in the rescue strain, confirming *fust-1’*s role in promoting circRNA production (Figure 1D). The *fust-1(csb21)* strain also showed another phenotype of lower average moving speed at day three adult stage when cultured at 25°C, which was also recovered in the rescue strain (Figure S1E). Although multiple copies of *fust-1* existed in the extrachromosomal arrays of the rescue and the overexpression strain (Figure S1F), these strains did not show much further improvement in circRNA levels or improvement in locomotion speed (Figure 1D and Figure S1E). This may be because of post-transcriptional regulation of *fust-1* or saturation of FUST-1 protein.

### 2. FUST-1 regulates circRNAs without affecting the cognate linear mRNAs

Next, to clarify whether FUST-1 promotes circRNA production by transcription promotion or not, levels of circRNAs and their cognate linear mRNAs were compared between the N2 strain and *fust-1(csb21)* strain at the L1 stage. While levels of these circRNAs were downregulated, their linear mRNA levels were not affected by the loss of FUST-1 (Figure 2A), indicating that FUST-1’s role in circRNA production is not through promoting transcription. Northern blot detection of the linear and circular transcript of *zip-2* also showed the same trend, in which circRNA was downregulated, whereas linear mRNA was not affected (Figure 2B and Figure 2C).

**Figure 2.**
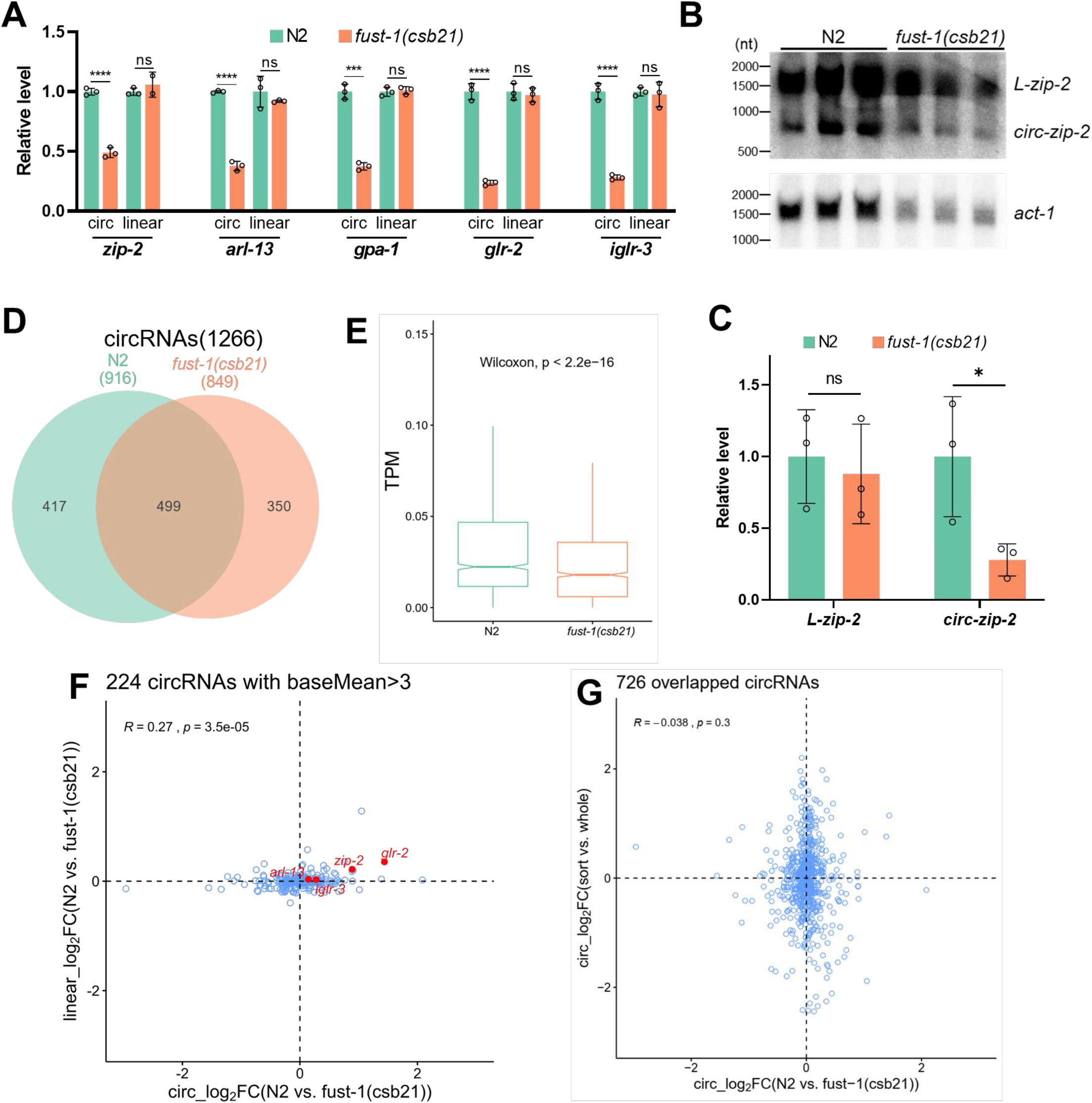
FUST-1 regulates circRNAs without affecting the cognate linear mRNAs. (A) RT-qPCR quantification of circRNAs and their linear mRNAs in the N2 strain and *fust-1(csb21)* strain. Levels are normalized to the N2 strain using *pmp-3* as the reference gene. Results are shown as mean ± sd of three biological replicates. Two-tailed Student’s *t*-test. \*\*\**p* < 0.001, \*\*\*\**p* < 0.0001; ns, not significant. (B) Northern blot detection *zip-2* transcripts, both linear and circular, and *act-1* in the L1 stage of N2 strain and *fust-1(csb21)* strain. (C) Quantification of northern blot results in (B), normalized to N2 strain using *act-1* as the reference gene. Results are shown as mean ± sd. Two-tailed Student’s *t*-test. \**p* < 0.05; ns, not significant. (D) Overlap of circRNAs detected in the RNA-seq results of N2 strain and *fust-1(csb21)* strain. (E) TPM (transcripts per million reads) comparison of all circRNAs between N2 and *fust-1(csb21). P* value indicates paired Wilcoxon test. (F) Scatter plot showing the log2 fold changes of 224 circRNAs with baseMean > 3 versus log2 fold changes of their corresponding linear mRNAs. The Pearson correlation coefficient (*R*) and *p* value (*p*) are shown. Names of several circRNA genes are labeled. (G) Scatter plot showing the log2 fold changes of 726 overlapped circRNAs between the “N2-*fust-1(csb21)*” dataset and the “sort-whole” dataset. The Pearson correlation coefficient (*R*) and *p* value (*p*) are shown.

To check the regulation of circRNA by FUST-1 globally, I performed RNA sequencing (RNA-seq) with ribosomal RNA depletion to compare differentially expressed circRNAs between *fust-1(csb21)* strain and wild-type N2 strain at the L1 stage. circRNA annotation was performed using DCC (Cheng et al., 2016), and differential expression was performed using DESeq2 (Love et al., 2014). Both mRNAs and circRNA clustered separately in principal component analysis (PCA) plots (Figure S2A and S2B). In total, 1266 circRNAs from 1199 annotated genes and 20 not-annotated loci were obtained with at least three back-splice junction (BSJ) reads in either group, with 916 in N2 strain and 849 in *fust-1(csb21)* strain (Figure 2D and Table S5). TPM (transcripts per million reads) values of circRNAs were compared between the two strains. circRNAs in the N2 strain showed significantly higher TPM values than those in *fust-1(csb21)* strain (paired Wilcoxon test, *p* < 2.2e-16) (Figure 2E), indicating general promotional roles of FUST-1 in circRNA production, although some circRNAs were also upregulated without FUST-1 (Table S6). Then, to check whether level changes in circRNA correlate with their cognate linear mRNAs, the fold changes of circRNA were plotted against those of their cognate mRNAs (Table S7), which showed a weak correlation (Pearson’s correlation coefficient *R* = 0.27, *p* = 3.5e-05) of circRNAs with baseMean (given by DESeq2) bigger than 3 (Figure 2F). The correlation was even weaker when all the circRNAs from annotated genes were considered (Figure S2C, Pearson’s correlation coefficient *R* = 0.14, *p* = 7.4e-07). These results were consistent with the finding that FUST-1 regulates circRNAs without disturbing the cognate linear mRNA levels (Figure 2A).

In my previous study, I provided the first neuronal circRNA profiles at the L1 stage of *C. elegans* (Cao, 2021). I then asked whether FUST-1 has a preference in the regulation of neuronal circRNAs. The circRNAs identified in this work (the “N2-*fust-1(csb21)*” dataset) were compared with my previous dataset (the “sort & whole” dataset), which resulted in 726 overlapped circRNAs (Figure S2D). Fold changes of the 726 overlapped circRNAs between N2 and *fust-1(csb21)* were plotted against those between the sort group (sorted neuron samples) and the whole group (whole worm samples), which showed no correlation (Figure 2G, Pearson’s correlation coefficient *R* = −0.038, *p* = 0.3), suggesting that FUST-1 has no preference for neuronal circRNAs.

### 3. FUST-1 binds to pre-mRNAs of circRNA genes

FUS binds to flanking introns of circRNA genes in N2a cells (Errichelli et al., 2017). I next checked whether FUST-1 in *C. elegans* recognizes pre-mRNAs of circRNA genes to regulate circRNA formation. Using CRISPR-Cas9 technology, I inserted the FLAG tag to the N terminal, just after the start codon, or to the C-terminal, just before the stop codon, respectively (Figure 3A and Figure S3A). The effect of FLAG-tag insertion on FUST-1’s role in circRNA regulation was evaluated. While N-terminal FLAG insertion showed slight increases in circRNA levels, C-terminal FLAG tag fusion affected FUST-1’s function in circRNA regulation in multiple circRNAs (Figure S3B and S3C). Hence N-terminal FLAG fused FUST-1 strain was used for the co-immunoprecipitation (Co-IP) experiment. Dynabeads Protein G conjugated with anti-FLAG antibody (+Ab) were used for Co-IP. Beads only (-Ab) were used as the negative control. As expected, the anti-FLAG antibody successfully enriched FLAG::FUST-1 after Co-IP (Figure 3B and Figure S3D). Then the levels of pre-mRNAs of circRNA genes were quantified by RT-qPCR. Threshold cycle (Ct) values were used for comparison. Lower Ct values indicate higher levels. While rRNA control (*18S rRNA* and *26S rRNA*) was depleted after Co-IP, all the pre-mRNAs of circRNA genes were enriched compared with input samples (Figure 3C). Moreover, these pre-mRNAs showed significate lower Ct values than those of control groups without using of antibody (Figure 3C), suggesting that FUST-1 binds to pre-mRNAs of the circRNAs genes to regulate circRNA formation.

**Figure 3.**
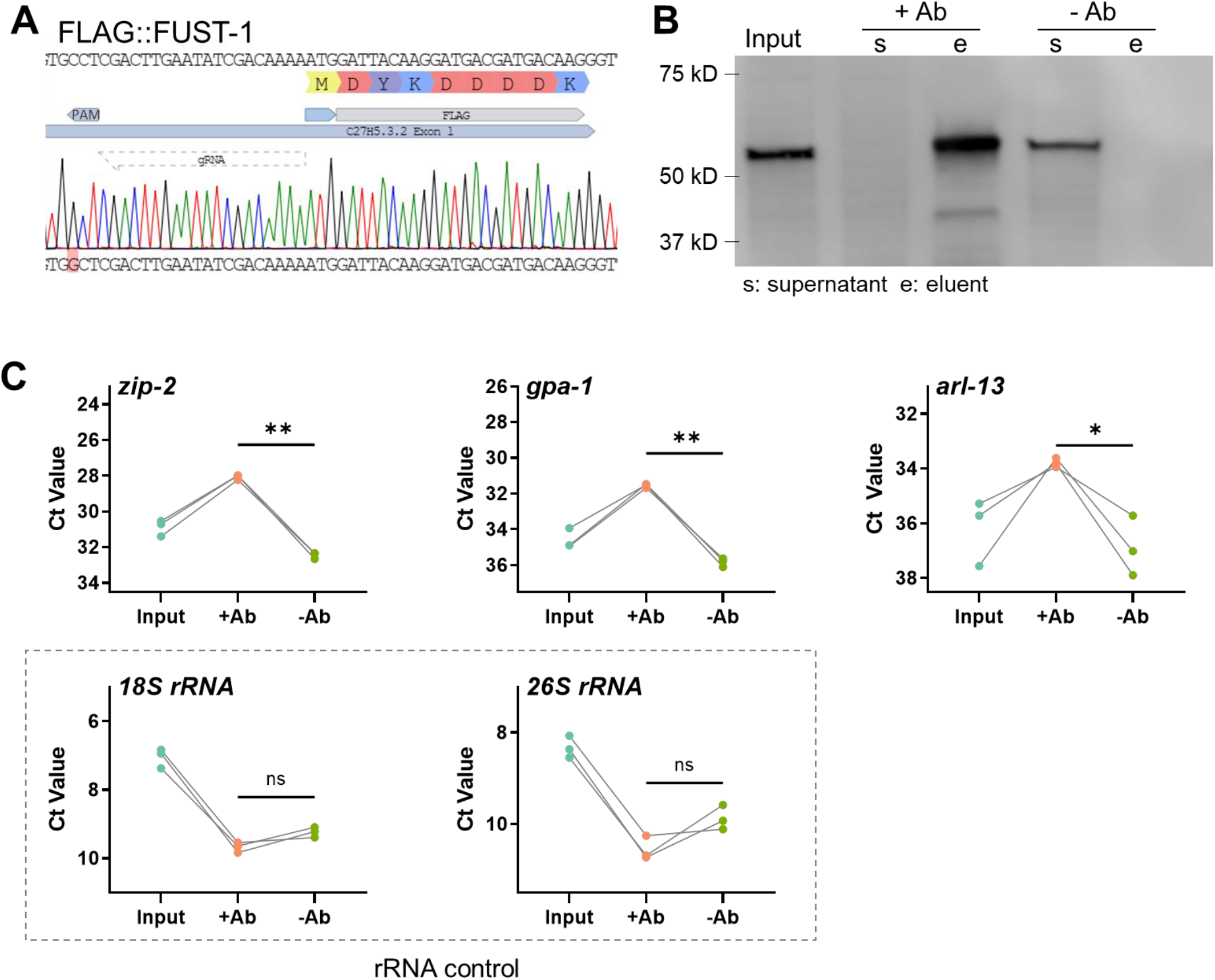
FUST-1 binds to pre-mRNAs of circRNA genes. (A) Sequence confirmation for N-terminal fusion of FLAG tag just after the start codon of FUST-1. Note the position of gRNA and the mutated PAM site (AGG>AGC). (B) Western blot showing the co-immunoprecipitation (Co-IP) of FLAG::FUST-1. (C) Ct value changes of pre-mRNAs of some circRNA genes before and after Co-IP of FLAG::FUST-1 with or without anti-FLAG antibody. Results from 3 biological replicates are shown. Paired two-tailed Student’s *t*-test. \**p* < 0.05, \*\**p* < 0.01, ns, not significant.

### 4. FUST-1 regulates both exon-skipping and back-splicing

In my previous study, transcripts that skip the exons to be circularized were identified in several circRNA genes (Cao, 2021). As the homolog of FUST-1 in humans and mice, FUS is involved in the regulation of AS of many genes by binding to their pre-mRNAs (Dichmann and Harland, 2012; Ishigaki et al., 2012; Rogelj et al., 2012). Since FUST-1 binds to the pre-mRNAs of these circRNA genes, I then checked whether FUST-1 could also regulate exon-skipping or not. In *zip-2*, reads aligned to the skipped junction were much less in *fust-1(csb21)* strain (27.0 reads on average) than those in wild-type N2 strain (71.3 reads on average) (Figure 4A). The RT-qPCR quantification results also showed that both the circRNA and the skipped transcript in *zip-2* were downregulated without FUST-1 (Figure 4B). In *arl-13*, while the circRNA got downregulated in *fust-1(csb21)*, the skipped transcript was weakly upregulated (Figure 4C). These results suggest that FUST-1 may function differently in different genetic environments.

**Figure 4.**
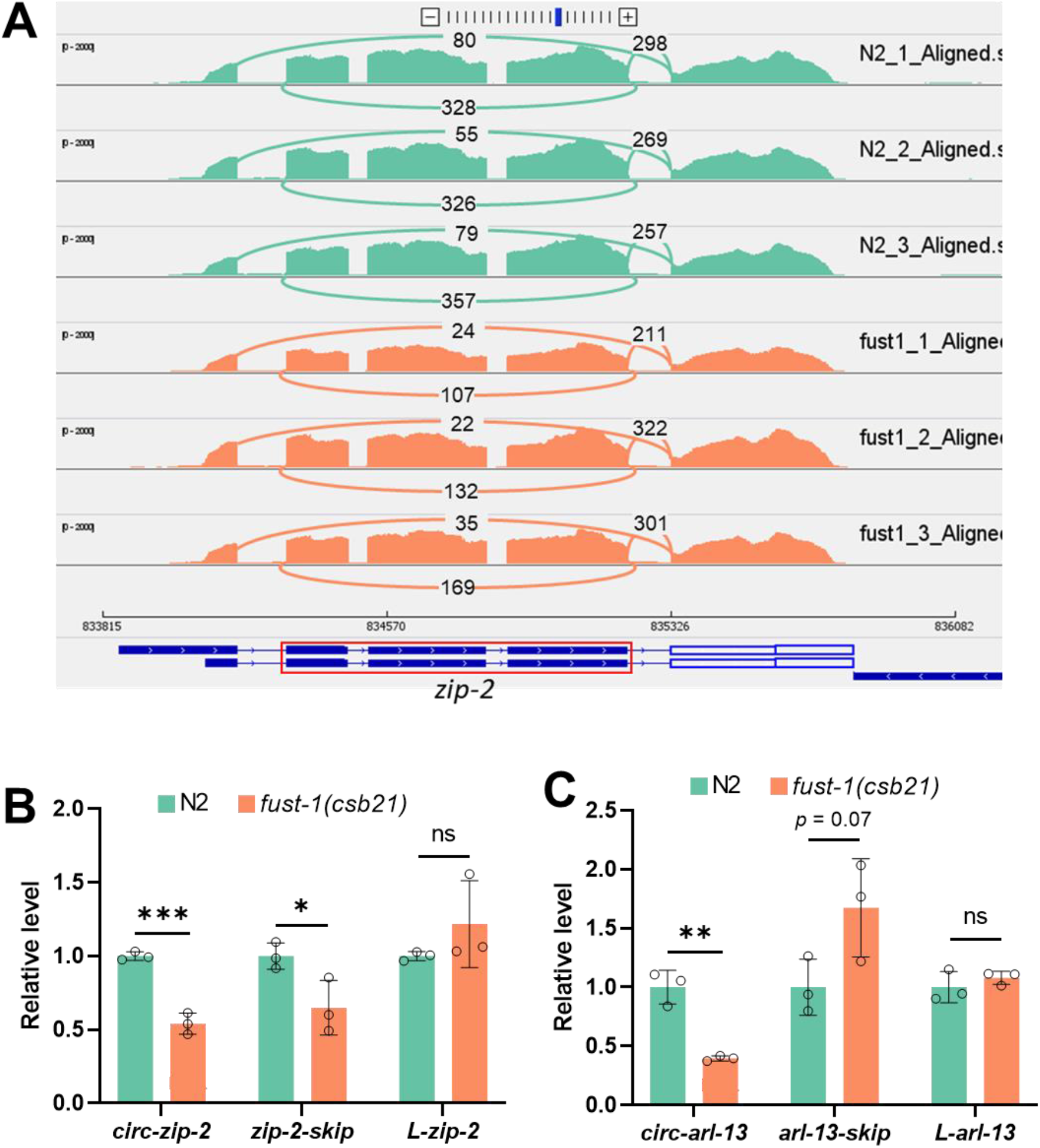
FUST-1 regulates both exon-skipping and back-splicing. (A) Sashimi plot showing numbers of reads aligned to the canonical splice junction, the skipped junction, and the back-splice junction in *zip-2*. Exons in the red rectangle are circularized. (B, C) RT-qPCR quantification of levels of the circular, skipped, and full-length linear transcripts in *zip-2* (B) and *arl-13* (C) between wild-type N2 strain and *fust-1(csb21)* strain. Levels are normalized to the N2 strain using *pmp-3* as the reference gene. Results are shown as mean ± sd of three biological replicates. Two-tailed Student’s *t*-test. \**p* < 0.05, \*\**p* < 0.01, *** *p* < 0.001, ns, not significant.

### 5. An autoregulation loop in *fust-1*

FUST-1 protein has two isoforms: isoform a is from the full-length transcript, and isoform b is from the transcript with skipped exon 5 (Figure 5A). Moreover, isoform b is translated using a downstream AUG and a different reading frame (+1) compared with isoform a. The reading frame in isoform b becomes the same as in isoform a after the skipping of exon 5 (38 nt in length). This results in a shorter isoform b with different N-terminal sequences, but the RNA recognition motif (RRM), zinc-finger (ZnF) domain, and the nuclear localization signal (NLS) domain are the same as isoform a (Figure 5A). To check how these two isoforms are expressed, two plasmids with different colors and a nonsense mutation in the reading frame of either isoform a (*fust-1a-mut::mRFP*) or isoform b (*fust-1b-mut::GFP*) were constructed so that only the other isoform can be expressed (Figure 5A and Figure S4A). Co-injection of the two plasmids in wild-type N2 strain showed that the two isoforms of FUST-1 were co-expressed in the nucleus of the same cells: neurons and intestinal cells (Figure 5B and Figure S4B). Interestingly, in early eggs, isoform a was expressed earlier than isoform b (Figure 5C). Furthermore, *fust1a-mut::GFP* plasmid expressed faintly in *fust-1(csb21)* strain and co-injection with *fust1b-mut::mRFP* can increase the GFP intensity (data not shown). These results gave a hint that isoform a may promote the production of isoform b.

**Figure 5.**
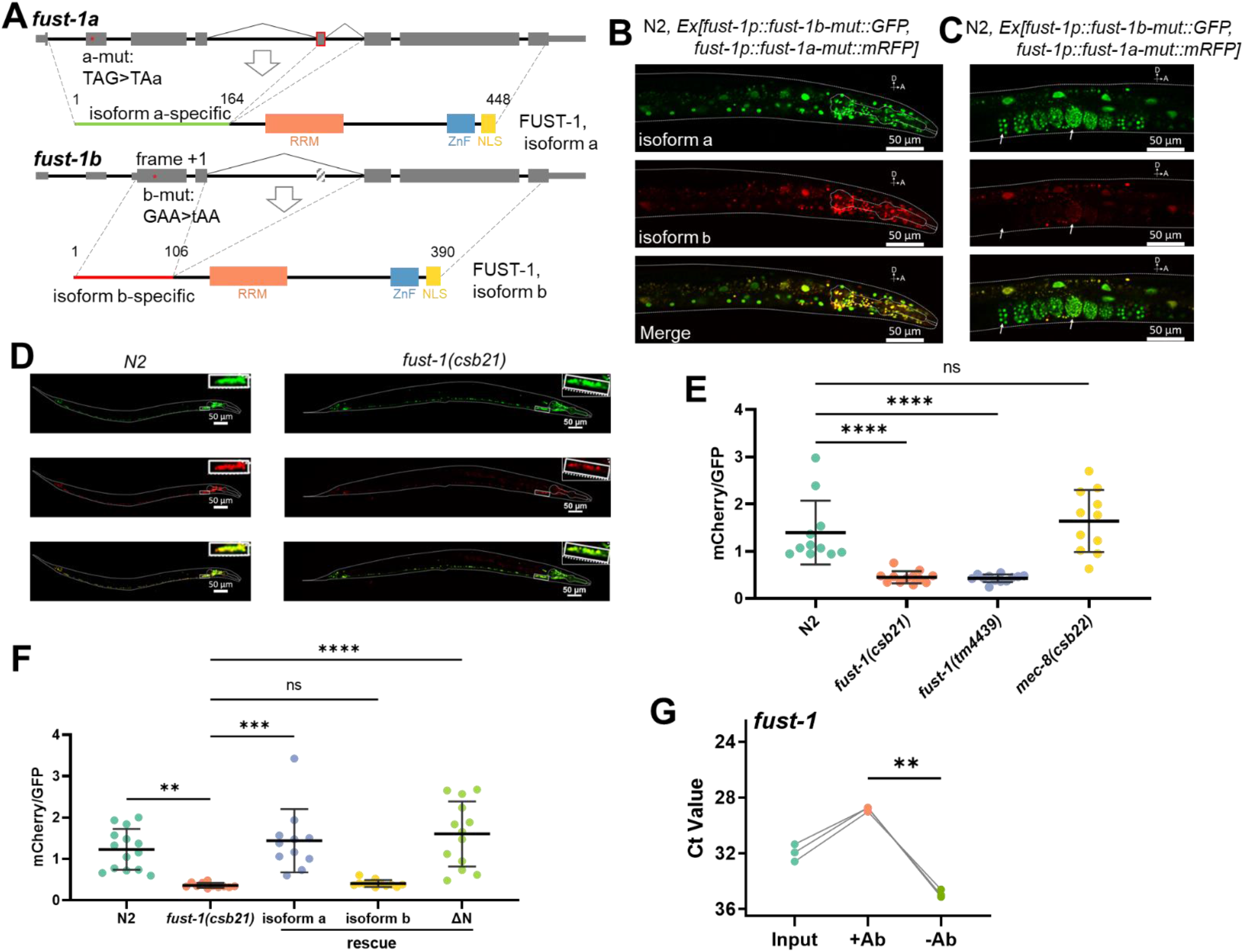
An autoregulation loop in *fust-1*. (A) Gene structure of *fust-1* and the domains in FUST-1 isoform a and isoform b. Note the positions where nonsense mutations were introduced. Lengths of amino acids in each isoform were labeled. RRM: RNA recognition motif; ZnF: Zinc-figure; NLS: nuclear localization signal. (B, C) Confocal images showing expression of FUST-1 isoform a and isoform b in the nucleus of neuron cells (B) and eggs (C). Note that in early eggs, FUST-1 isoform a was expressed earlier than isoform b (white arrows). A: Anterior, D: Dorsal. Scale bars: 50 μm. (D) Representative confocal images showing the expression patterns of splicing reporter of *fust-1* exon 5 in N2 strain and *fust-1(csb21)* strain. Inset squares show the enlarged neck neurons in indicated strains. A: Anterior, D: Dorsal. Scale bars: 50 μm. (E, F) Quantification of mCherry-to-GFP ratios of the *fust-1* exon5 splicing reporter in the indicated strains. One-way ANOVA, Tukey’s multiple comparisons. \*\**p* < 0.01, \*\*\**p* < 0.001, \*\*\*\**p* < 0.0001; ns, not significant. (G) Ct value changes of *fust-1* pre-mRNA before and after Co-IP of FLAG::FUST-1 with or without anti-FLAG antibody. Primer positions are in intron 4 of *fust-1* pre-mRNA. Results from 3 biological replicates are shown. Paired two-tailed Student’s *t*-test. \*\**p* < 0.01.

To prove this hypothesis, I constructed a dual-color splicing reporter (Norris et al., 2014; Thompson et al., 2019) of the skipping of exon 5 in *fust-1* (Figure S4C), in which no skipping gives GFP expression while skipping of exon 5 results in mCherry expression. As expected, two colors were co-expressed in almost all the neurons in wild-type strain (Figure 5D), suggesting that exon-skipping of exon 5 is happening in all the neurons. However, when the reporter plasmid was crossed into two *fust-1* mutation strains, *fust-1(csb21)* and *fust-1(tm4439)* (Figure S1C), the expression of mCherry was dramatically reduced (Figure 5D and Figure S4D), indicating FUST-1 was involved in the exon-skipping of its own pre-mRNA. Since *fust-1(csb21)* strain has pharyngeal GFP expression (Norris et al., 2017) (Figure S1C and Figure 5D), neurons in the ventral nerve cord around the neck were used to quantify the mCherry-to-GFP intensity ratios (Figure 5D). The mCherry-to-GFP ratios were significantly reduced in both two *fust-1* mutants, and they did not change in the *mec-8(csb22)* strain (Figure 5E and Figure S4D).

Next, to prove that FUST-1, isoform a promotes the skipping of exon 5 of *fust-1* pre-mRNA, I tried the rescue of mCherry expression of the splicing reporter in *fust-1(csb21)* by co-injection of the reporter plasmid with FUST-1, isoform a cDNA or FUST-1, isoform b cDNA, driven by the *fust-1* original promoter (2181 bp upstream the ATG of FUST-1, isoform a). One more construct with truncated N-terminal (FUST-1, ΔN) was also used (Figure S5A). Tail-expressing plasmid *lin-44p::mRFP* was used as an injection marker. As expected, isoform a cDNA restored the mCherry expression of the splicing reporter, while isoform b cDNA did not (Figure 5F, Figure S5B and S5C), which confirms that FUST-1, isoform a promotes the skipping of exon-5 to produce FUST-1 isoform b. Consistent with this, *fust-1* pre-mRNA, detected by primers in intron 4 of *fust-1*, was significantly enriched after Co-IP with FLAG::FUST-1, which only tagged FUST-1, isoform a (Figure 5G). To my surprise, the FUST-1, ΔN construct also rescued the mCherry expression, just as efficient as FUST-1, isoform a (Figure 5F and Figure S5D). Since the three isoforms have identical functional domains (RRM, ZnF, and NLS) with different N-terminal sequences, these results suggest that the N-terminal in FUST-1, isoform b may prevent its domains from functioning normally. Taken together, I characterized an autoregulation loop in *fust-1*, in which FUST-1, isoform a promote the skipping of exon 5 of *fust-1* pre-mRNA, resulting in the production of FUST-1, isoform b.

### 6. FUST-1, isoform a is the functional isoform in circRNA regulation

Next, to check which isoform of FUST-1 is functional in circRNA regulation, I tried to rescue the downregulated circRNAs in *fust-1(csb21)* with extrachromosomal expression of FUST-1 isoform cDNA with C-terminal mRFP fusion, in which either FUST-1, isoform a, or FUST-1, isoform b or FUST-1, ΔN is expressed (Figure S6A). The mRFP-positive L1 worms were sorted, from which total RNA was extracted, and then circRNA levels were quantified by RT-qPCR. Same with their roles in exon-skipping, FUST-1, isoform a successfully rescued the downregulated circRNAs, whereas FUST-1, isoform b did not improve the downregulated circRNA levels at all, indicating that FUST-1, isoform a is the functional protein in circRNA regulation (Figure 6A). Although not as efficient as FUST-1, isoform a, FUST-1, ΔN fully rescued the downregulated *circ-zip-2* and *circ-iglr-3* and partially restored *circ-arl-13* level (Figure 6A).

**Figure 6.**
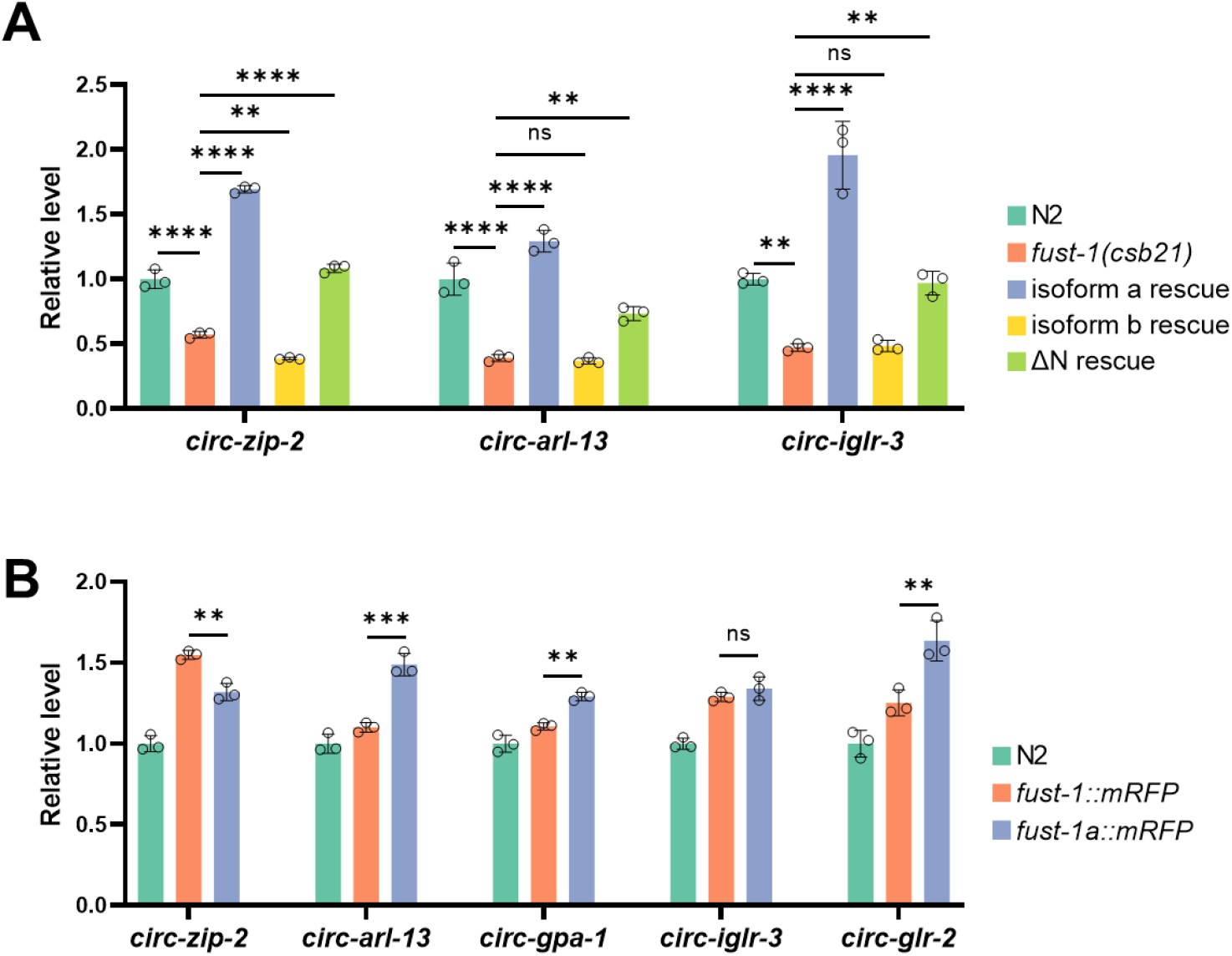
FUST-1, isoform a is the functional isoform in circRNA regulation. (A) Rescue of circRNA levels by FUST-1 isoforms, quantified by RT-qPCR. cDNA samples from L1 worms of indicated strains were used. (B) RT-qPCR quantification of circRNA levels at the L1 stage of indicated strains. (A, B) Levels are normalized to N2 strain using *pmp-3* as the reference gene. Results are shown as mean ± sd of three biological replicates. One-way ANOVA, Tukey’s multiple comparisons. \*\**p* < 0.01, \*\*\**p* < 0.001, \*\*\*\**p* < 0.0001; ns, not significant.

In an effort to generate strains with C-terminal mRFP tagging of FUST-1 isoforms, I achieved C-terminal mRFP insertion in *fust-1 (fust-1::mRFP)*. Another obtained strain, in which intron 3 to intron 6 of *fust-1* were removed, cannot use the autoregulation pathway, resulting in the expression of only FUST-1, isoform a (*fust-1a::mRFP*) (Figure S6B). I failed to obtain a strain that can only express mRFP tagged FUST-1, isoform b. Consistent with the extrachromosomal expression pattern of FUST-1 (Figure 1C and Figure S1D), mRFP-tagged FUST-1 was mainly expressed in the nucleus of neurons and intestinal cells (Figure S6C). Moreover, FUST-1 was also found in the nucleus of gonads (Figure S6D), which was not observed in extrachromosomal expression, probably due to silencing of the multicopy transgenes in the germline (Merritt and Seydoux, 2010).

Levels of circRNAs were compared between the two strains. Out of the five checked circRNA, the levels of four circRNAs were altered in the strain only FUST-1, isoform a can be expressed (Figure 6B), suggesting the autoregulation loop is critical for FUST-1’s role in circRNA regulation.

## Discussion

Using identified circRNAs in the neurons as targets, I performed a small-scale screening of thirteen conserved RBP genes in their roles of circRNA regulation. Most of these RBPs showed promotional roles in circRNA production, suggesting that the involvement of RBPs in back-splicing may be common in *C. elegans*. I further showed that FUST-1, the homolog gene of FUS in *C. elegans*, regulates circRNA formation without affecting the cognate linear mRNAs (Figure 2). Although I used circRNAs either enriched in neurons or highly expressed in neurons as targets to identify FUST-1, FUST-1 did not show preference in the regulation of neuronal circRNAs (Figure 2G). Since FUST-1 is also expressed in intestine and germline cells, FUST-1 may regulate circRNAs in those cells.

CLIP-seq data on FUS suggest that rather than recognizing specific sequences, FUS tends to bind to stem-loop secondary structures (Hoell et al., 2011; Ishigaki et al., 2012; Rogelj et al., 2012; Zhou et al., 2013). In cultured N2a cells, FUS binds to the flanking introns of circularized exons (Errichelli et al., 2017). In this study, I found that FUST-1 binds to the pre-mRNAs of circRNA genes. Interestingly, I found that FUST-1 regulates both back-splicing and exon-skipping in *zip-2* and *arl-13*. In my previous work, I discovered that RCMs in circRNA-flanking introns of *zip-2* simultaneously promote both exon-skipping and back-splicing (Cao, 2021). It is possible that once FUST-1 recognizes sequences in circRNA-flanking introns in *zip-2*, it can regulate both processes at the same time.

Self-regulation has been reported in FUS, where FUS promotes skipping of exon 7 of its pre-mRNA, which results in NMD (Zhou et al., 2013). Unlike the previous example, FUST-1, isoform a-promoted exon skipping of *fust-1* pre-mRNA produces FUST-1, isoform b that contains exactly the same functional domains, but with different N-terminal sequences. While FUST-1, isoform a is capable of promoting exon-skipping and circRNA regulation, FUST-1, isoform b is not functional in either of the two aspects (Figure 5F and Figure 6A). I first hypothesized that N-terminal sequences in FUST-1, isoform a may be important for its function. However, the FUST-1, ΔN construct, which has no N-terminal sequences, seemed functional in both exon-skipping promotion and circRNA regulation, although not as efficient as FUST-1, isoform a. These results suggest that N-terminal in isoform b may interfere with the functional domain(s), possibly RRM, so that FUST-1, isoform b cannot bind to the target mRNAs recognized by FUST-1, isoform a.

The frameshifting in FUST-1, isoform b dramatically changes the N-terminal amino acid contents compared with FUST-1, isoform a. The isoform a-specific N-terminal has high ratios of glycines (53/164, 32.3%) and glutamines (22/164, 13.4%). However, isoform b-specific N-terminal contains more valines (18/106, 17.0%) and glutamic acid residues (18/106, 17.0%), which are very few in isoform a-specific N-terminal: 0/164 and 2/164, respectively. High valine content may make the N-terminal of FUST-1, isoform b more hydrophobic and high glutamic acid content can add more negative charges, which may cause the folding of FUST-1, isoform b different from isoform a. Further *in vitro* RNA binding experiments or structural analysis may be worth trying to investigate the detailed mechanisms that dictate different function potentials in the two FUST-1 isoforms.

## Acknowledgment

I thank Dr. Adam Norris from the Department of Biological Sciences at Southern Methodist University for providing the RBP strains and the dual-color splicing reporter plasmids. I thank Dr. Alex Parker from the Department of neuroscience, Université de Montréal for providing the *fust-1(tm4439)* strain. I thank DNA Sequencing Section of OIST for its help on library preparation and RNA sequencing of N2 and *fust-1(csb21)* samples. I thank the Information Processing Biology Unit (Maruyama Unit) members for their discussion and feedback. I am grateful for the help and support provided by the Scientific Computing and Data Analysis section of the Research Support Division at OIST. I thank Okinawa Institute of Science and Technology, Graduate School for financial support.

## Author Contributions

D.C. designed and conducted all the experiments, performed all the analysis, and wrote the paper.

## Materials and methods

### Worm maintenance

*C. elegans* Bristol N2 strain was used as the wild type. Worms were maintained using standard conditions on Nematode Growth Media (NGM) agar plates with *Escherichia coli* strain OP50 (Brenner, 1974) at 20°C or 25°C. New transgenic worms were generated by microinjection with ~40 ng/μl plasmid. The strains used in this study are listed in TableS1.

### Plasmid preparation

*fust-1p::fust-1::mRFP: fust-1* genomic fragment containing sequences from 2181bp upstream ATG to just before stop codon was cloned into the Sma I site of pHK-mRFP vector in frame with mRFP by In-Fusion HD Cloning Kit (Takara). This plasmid was further used to generate backbone structure containing *fust-1* promoter and mRFP, to which cDNAs of FUST-1 isoforms (isoform a, isoform b, and ΔN) were inserted by In-Fusion (Takara). The mRFP fused FUST-1 cDNA plasmids were used to generate cDNA only plasmids for splicing reporter rescue by removing the mRFP sequences using In-Fusion (Takara). Splicing reporter of *fust-1* exon 5 was prepared by cloning exon 4 to exon 6 into the plasmids provided by Dr. Adam Norris.

### Worm synchronization

Worm synchronization was performed by bleaching for large-scale worm preparation (RNA extraction). For small-scale worm preparation (locomotion assay), worms were synchronized by egg-laying. Briefly, 10 - 15 gravid adult worms were placed onto a seeded NGM plate for four hours and worms were removed after egg-laying. The eggs were then cultured to the desired stage.

### Worm sorting

L1 worms with extrachromosomal fluorescent proteins were obtained by bleaching and hatching overnight at room temperature. Then fluorescence-positive worms were sorted using BioSorter Large Particle Flow Cytometer (Union Biometrica).

### Mutagenesis by CRISPR-Cas9

Mutation by CRISPR-Cas9 was performed as described previously (Cao, 2021). To make the recombinant dsDNAfragments for insertion, plasmids containing the wild-type FUST-1::mRFP were constructed first, from which dsDNA repair fragments were amplified using primers containing recombinant sequences. Guide RNA sequences, recombinant ssDNAs, validation primers, and primers for recombinant fragment amplification are listed in Table S3.

### Co-immunoprecipitation (Co-IP)

~20,000 L1 FLAG::FUST-1 worms were seeded on a nutrition enriched plate with NA22 *E. coli* (NEP-NA22). After 4-day culture at 20°C, all bacteria were consumed, and most of the progenies were at the L1 stage. Adult worms were removed by filtering through a 30 μm mesh. Three NEP-NA22 plates, which gave ~ 1 million L1 worms, were used for one replicate experiment. Worms were washed with 1 × 10 ml M9 buffer, 2 × 10 ml cold Buffer B70 (50 mM HEPES-KOH (pH 7.4), 70 mM potassium acetate (KAc), 1 mM sodium fluoride (NaF), 20 mM β-glycerophosphate, 5 mM magnesium acetate (MgOAc), 0.1% Triton X-100, 10% glycerol). Worms were then resuspended in 0.4 ml Buffer B70 supplemented with 2 × cOmplete Proteinase inhibitor cocktail (Roche) and dripped into liquid N2 with 1 ml pipette tips to form small pearls. Worm pearls were stored at −80°C. Worm pearls were ground into fine powder in a mortar containing liquid N2, which was suspended into 1 ml cold Buffer B70 supplemented with 2 × cOmplete Proteinase inhibitor cocktail (Roche) and 5 μl Murine RNase Inhibitor (NEB). Worm lysate was cleared by centrifugation at 20,000 × g for 20 min at 4°C. 50 μl worm lysate was taken as input samples, in which 40 μl was used for RNA extraction and 10 μl for western blot. 50 μl Dynabeads Protein G (Invitrogen) was coupled with or without 5 μg Anti-FLAG M2 antibody (Sigma-Aldrich), which was then incubated with 400 μl lysate, rotating overnight at 4°C. The next day, the lysate-beads slurry was cleared magnetically, and the supernatant was taken for western blot. Keeping tubes on magnetic tray, the beads were washed 2 × 200 ul Buffer B70 gently. 50 μl 50 mM glycine, pH 2.8 was added to the washed beads to elute bound RBP complex. After mixing and incubating at RT for 3 min, the supernatant was transferred to another tube containing 5 μl 1 M Tris-HCl, pH 7.5 for pH neutralization. For the 55 μl elution, 44 μl was used for RNA extraction, 11 μl for western blot.

### Western blot

Protein samples were resolved by SDS-PAGE (5% stacking gel and 12% resolving gel) and transferred to PVDF membrane by the standard protocol (25 V, 30 min) of Trans-Blot Turbo Transfer System (Bio-Rad). After blocking with 5% BSA-PBST (137 mM Sodium Chloride, 10 mM Phosphate, 2.7 mM Potassium Chloride, pH 7.4, 0.1% (v/v) Tween-20, and 5% (w/v) BSA) for 1 hour at room temperature, the membrane was incubated overnight with primary antibody (listed below) at 4°C. After 3 × 5 min washes in PBST, the membrane was incubated with HRP-conjugated secondary antibody at room temperature for 1 hour. The membrane was washed 3 × 5 min in PBST and then visualized by Amersham ECL Prime Western Blot Detection Reagent (GE Healthcare). Images were taken by Fluorescent Image Analyzer LAS-3000 (FujiFilm) using the chemiluminescence channel. Mouse ANTI-FLAG M2 antibody (F3165, Sigma-Aldrich): 1:2000; Amersham ECL Mouse IgG, HRP-linked whole Ab (from sheep):1:2000.

### RNA extraction

RNA extraction was performed using Direct-zol RNA MicroPrep kit (ZYMO Research) with on-column DNase I (ZYMO Research) digestion according to the manufacturer’s protocol. For RNA extraction from worms, worms were first flash-frozen in Trizol solution (Invitrogen) in liquid N2 and then homogenized by vortexing with glass beads (φ 0.1 mm) in Beads Cell Disrupter MS-100 (TOMY).

### RNA Sequencing

For RNA-seq of samples from the L1 stage of N2 and *fust-1(csb21)*, 500 ng total RNA samples from 3 biological duplicates were used as inputs. rRNA depletion was performed using Ribo-Zero Plus rRNA Depletion kit (Illumina) and library preparation was conducted using NEBNext Ultra II Directional RNA Library Prep Kit for Illumina (New England BioLabs) according to manufacturer’s protocols. Sequencing was performed on NovaSeq 6000 (Illumina) to obtain 150 nt paired-end reads.

### Real-time PCR

Real-time PCR reactions were performed using soAdvanced Universal SYBR Green Supermix (Bio-Rad) with cDNAs synthesized from iScript Advanced cDNA synthesis kit (Bio-Rad). 20 μl reaction mix with 2 μl cDNA (~1-10 ng) were monitored on StepOnePlus Thermal Cycler (Applied Biosystems) in “fast mode”. Cycling conditions: 95 °C, 30’, 40 or 45 cycles of 95 °C, 15’ and 60 °C, 30’with plate reading, and a final melt curve stage using default conditions. If not mentioned, all cDNAs used for RT-qPCR were from the L1 stage of indicated strains. If Ct values were not determined or higher than Ct values in no-template control (NTC) samples, they are treated as “n.d.” (not detected). Primers used for RT-qPCR are listed in Table S2.

### Northern blot

Northern blot was performed using NorthernMax kit (ThermoFisher Scientific) as described previously (Cao, 2021). Primers used for probe amplification are in Table S2.

### circRNA prediction and RNA-seq data analysis

circRNA analysis and differential expression analysis were performed as described previously (Cao, 2021). In the dataset in this study, many circRNAs were predicted from *rrn-3.1*, which were removed for downstream analysis. The plots (PCA plots, boxplots, scatter plots) were generated using ggplot2 package (https://ggplot2.tidyverse.org/), and ggpubr (http://www.sthda.com/english/rpkgs/ggpubr) package in R.

### Microscopy

Confocal images were obtained using a Zeiss LSM780 confocal microscope. For mCherry-to-GFP ratio quantification of splicing reporter of *fust-1* exon 5, all images were taken under the same setting parameters (Pinhole: 1.00 AU; Laser: 561 nm, 2.00%, 488 nm, 2.00%; Detection wavelength: GFP, 493-556 nm, mCherry, 588-694 nm; Gain: GFP, 625.0, mCherry, 790.0; Detector Digital Gain: 1.0 for all channels) to make sure that no saturation in both GFP and mCherry channels. Images were processed using ZEISS ZEN3.1 software. The average intensities in the GFP channel and the mCherry channel were used for quantification.

### Locomotion Assay

Locomotion analysis of day 3 adult worms was performed as described previously (Kawamura and Maruyama, 2019). Briefly, 15 synchronized day 3 adult worms were picked onto a blank NGM plate to get rid of food for ~1 min. The worms were then transferred to another empty NGM plate and locomotion images were recorded for 1min with five frames with the lid on. Images were analyzed using ImageJ and wrMTrck plugin (Nussbaum-Krammer et al., 2015) (http://www.phage.dk/plugins/wrmtrck.html) to calculate the average speeds. More than 50 worms were recorded. Worms that got lost during recording were not included.

### Data availability

Raw FASTQ files from the RNA-seq data were deposited at the NCBI Sequence Read Archive (BioProject: PRJNA669975, Table S4). All strains, plasmids, and other materials are available upon request.

---

**Figure S1:**
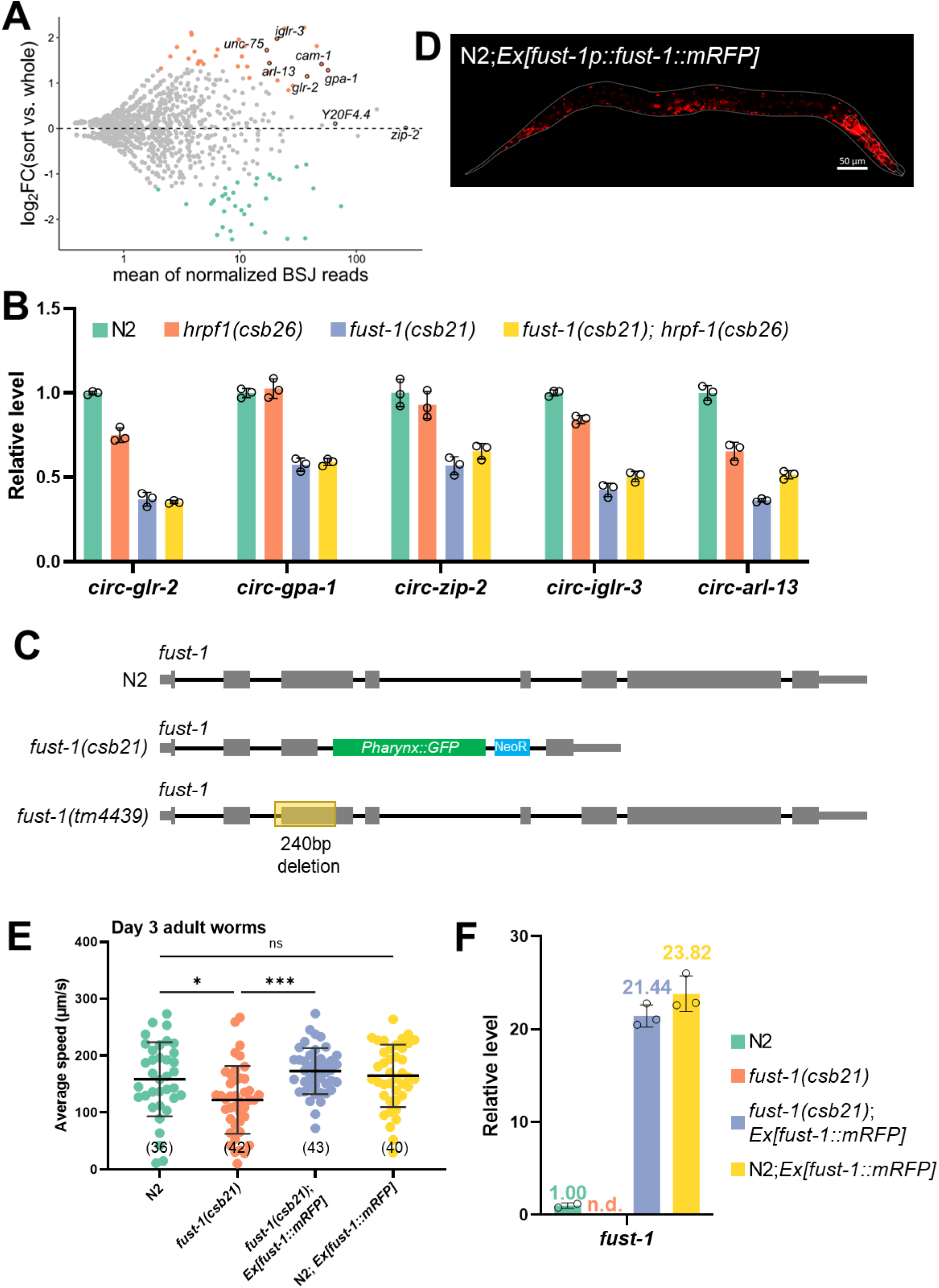
Related to Figure 1. (A) Differentially expressed circRNAs between sorted neuron samples and whole worm samples. Colored dots represent significantly differentially expressed circRNAs. (B) RT-qPCR quantification of circRNA levels at L1 stage of indicated strains. Note that there is no additive effect in *fust-1(csb21); hrpf-1(csb26)* double mutant strain compared with *fust-1(csb21)* strain. Levels are normalized to the N2 strain using *pmp-3* as the reference gene. Results are shown as mean ± sd of three biological replicates. (C) Gene structure of *fust-1* in wildtype N2 strain, *fust-1(csb21)*, and *fust-1(tm4439)* strain. (D) Representative image showing mRFP-fused FUST-1 in wildtype N2 strain. Scale bar: 50 μm. (E) Average moving speed of day 3 adult worms raised at 25°C with the indicated genotypes. Numbers in brackets are numbers of worms used for moving speed measurement. Results are shown as mean ± sd. One-way ANOVA, Tukey’s multiple comparisons. \**p*<0.05, \*\*\**p* < 0.001, ns, not significant. (F) RT-qPCR quantification of fust-1 mRNA levels at the L1 stage of indicated strains. Levels are normalized to the N2 strain using *pmp-3* as the reference gene. Results are shown as mean ± sd of three biological replicates. n.d.: not detected.

**Figure S2:**
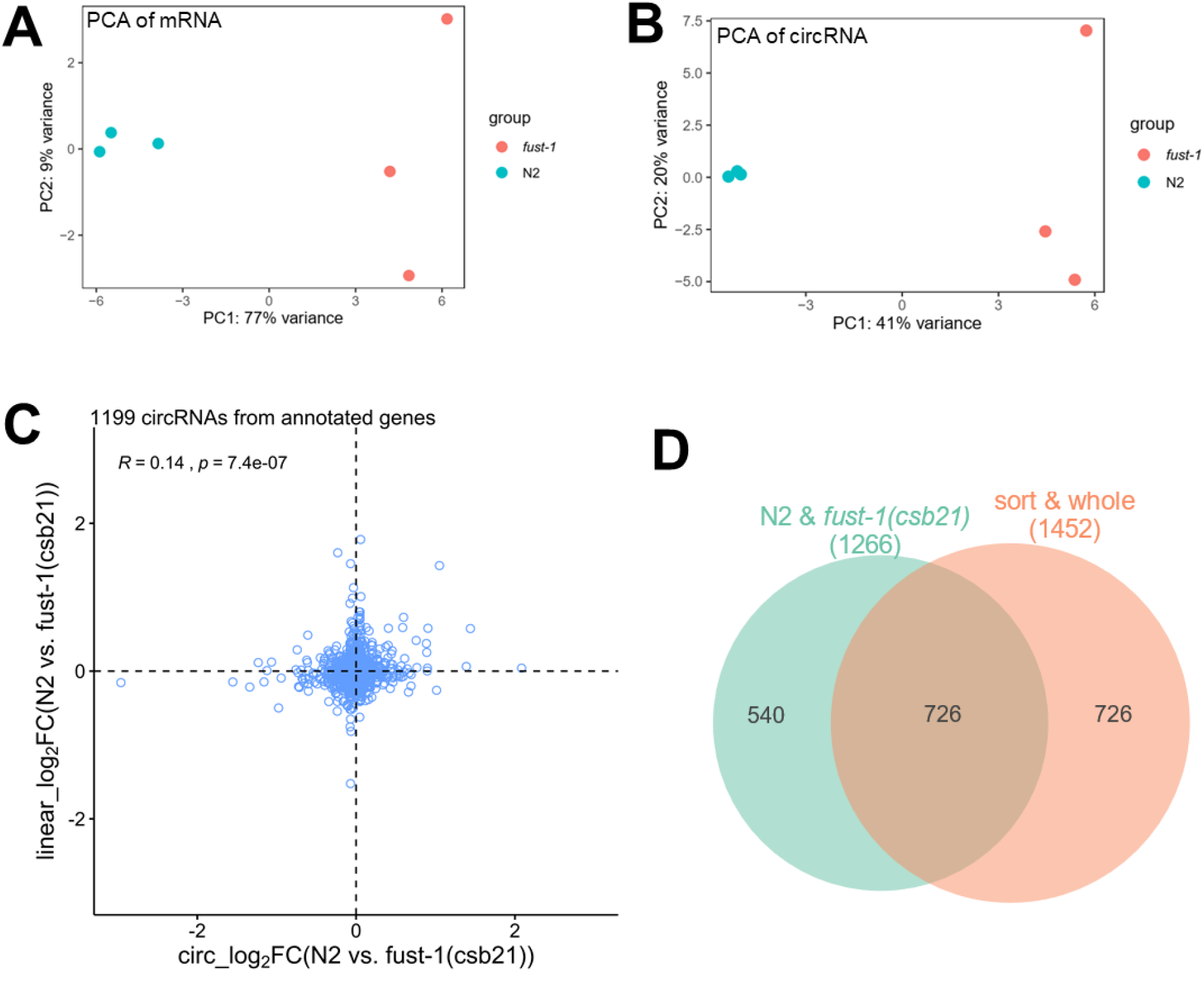
related to Figure 2. (A, B) Principal component analysis of linear mRNAs (A) and circRNAs (B) between wildtype N2 strain and *fust-1(csb21)* strain. (C) Scatter plot showing the log2 fold changes of all circRNAs versus log2 fold changes of their cognate linear mRNAs. The Pearson correlation coefficient (*R*) and *p* value (*p*) are shown. (D) Venn diagram showing overlapped circRNAs between the “N2-*fust-1(csb21)*” dataset and the “sort & whole” dataset.

**Figure S3:**
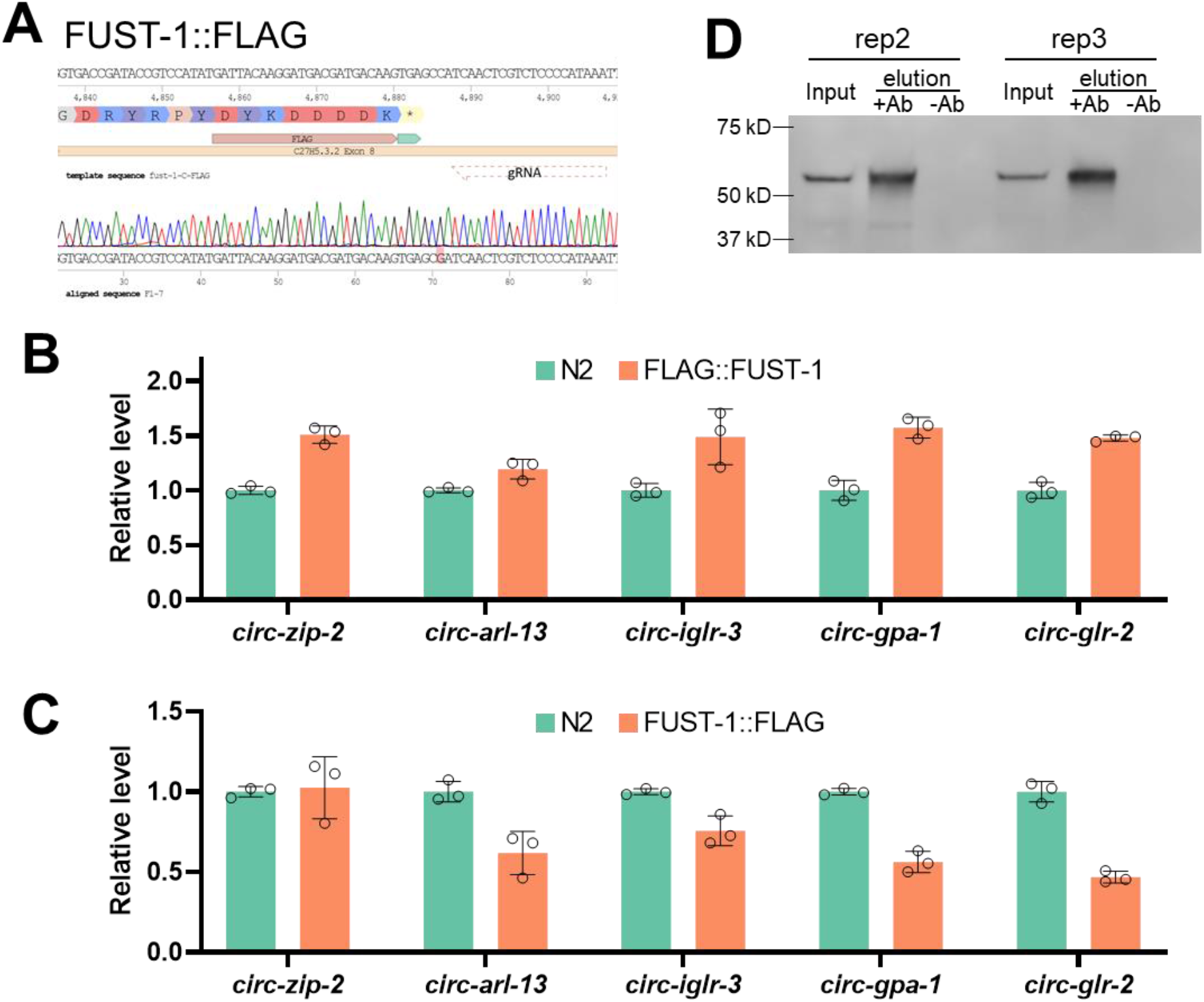
related to Figure 3. (A) Sequence confirmation of C-terminal fusion of FLAG tag just before the stop codon of FUST-1. Note the position of gRNA and the mutated PAM site (TGG>TCG). (B, C) RT-qPCR quantification of circRNA levels between wildtype N2 strain and N-terminal FLAG fusion of FUST-1 strain (B) and between wildtype N2 strain and C-terminal FLAG fusion of FUST-1 strain (C). Levels are normalized to the N2 strain using *pmp-3* as the reference gene. Results are shown as mean ± sd of three biological replicates. (D) Western blot of the other two biological replicates of FLAG::FUST-1 Co-IP with or without FLAG antibody.

**Figure S4:**
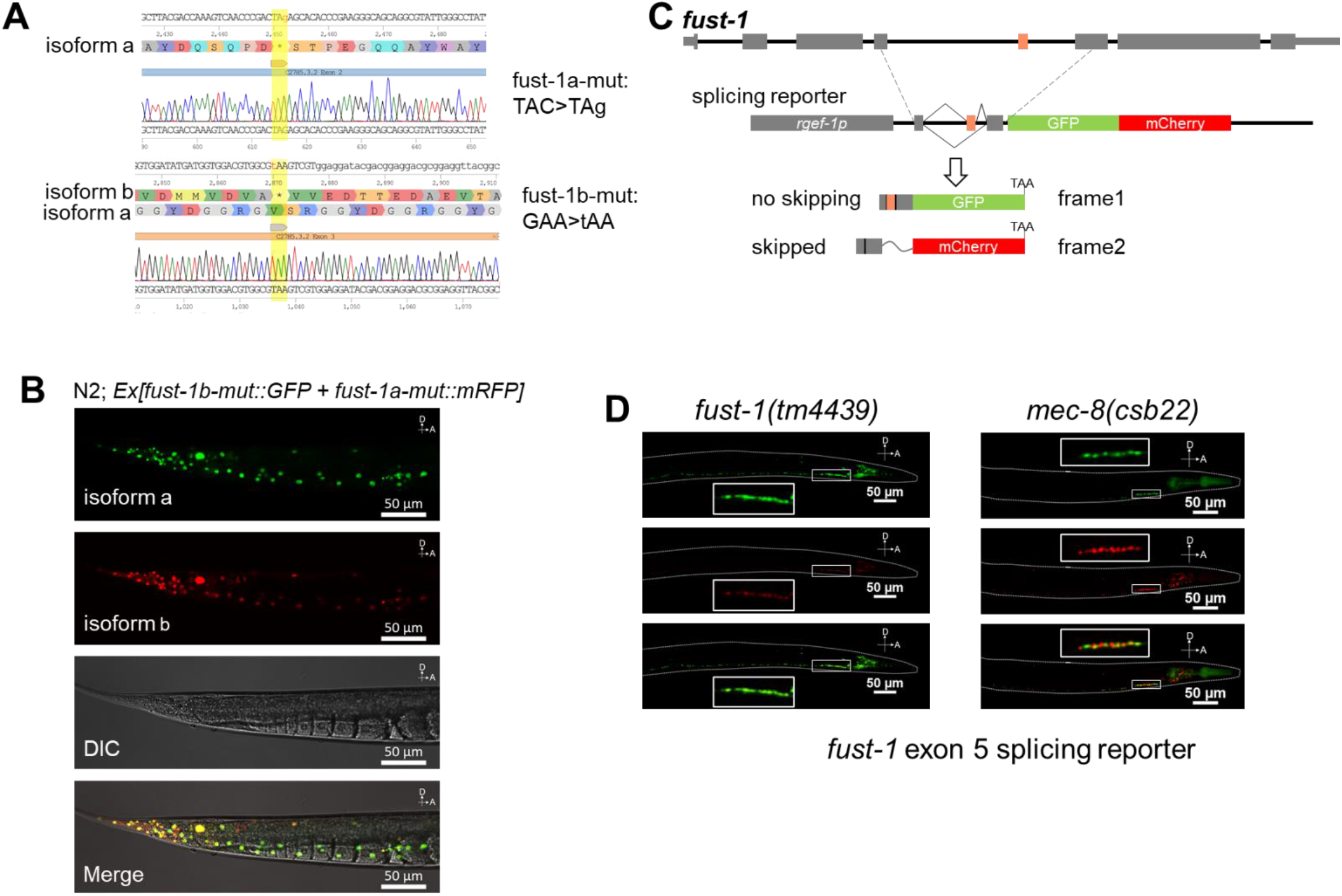
related to Figure 5. (A) Sequence confirmation of *fust-1a-mut::mRFP* plasmid and *fust-1b-mut::GFP* plasmid. The mutated sites are in yellow shadows. Note the introduction of the stop codons in the read frame of isoform a and isoform b. (B) Confocal images showing expression of FUST-1 isoform a and isoform b in the tail. A: Anterior, D: Dorsal. Scale bars: 50 μm. (C) Illustration of the dual-color splicing reporter of *fust-1* exon 5. (D) Representative images showing the expression patterns of splicing reporter of *fust-1* exon 5 in *fust-1(csb21)* strain and *fust-1(tm4439)* strain. Inset squares show the enlarged neck neurons. A: Anterior, D: Dorsal. Scale bars: 50 μm.

**Figure S5:**
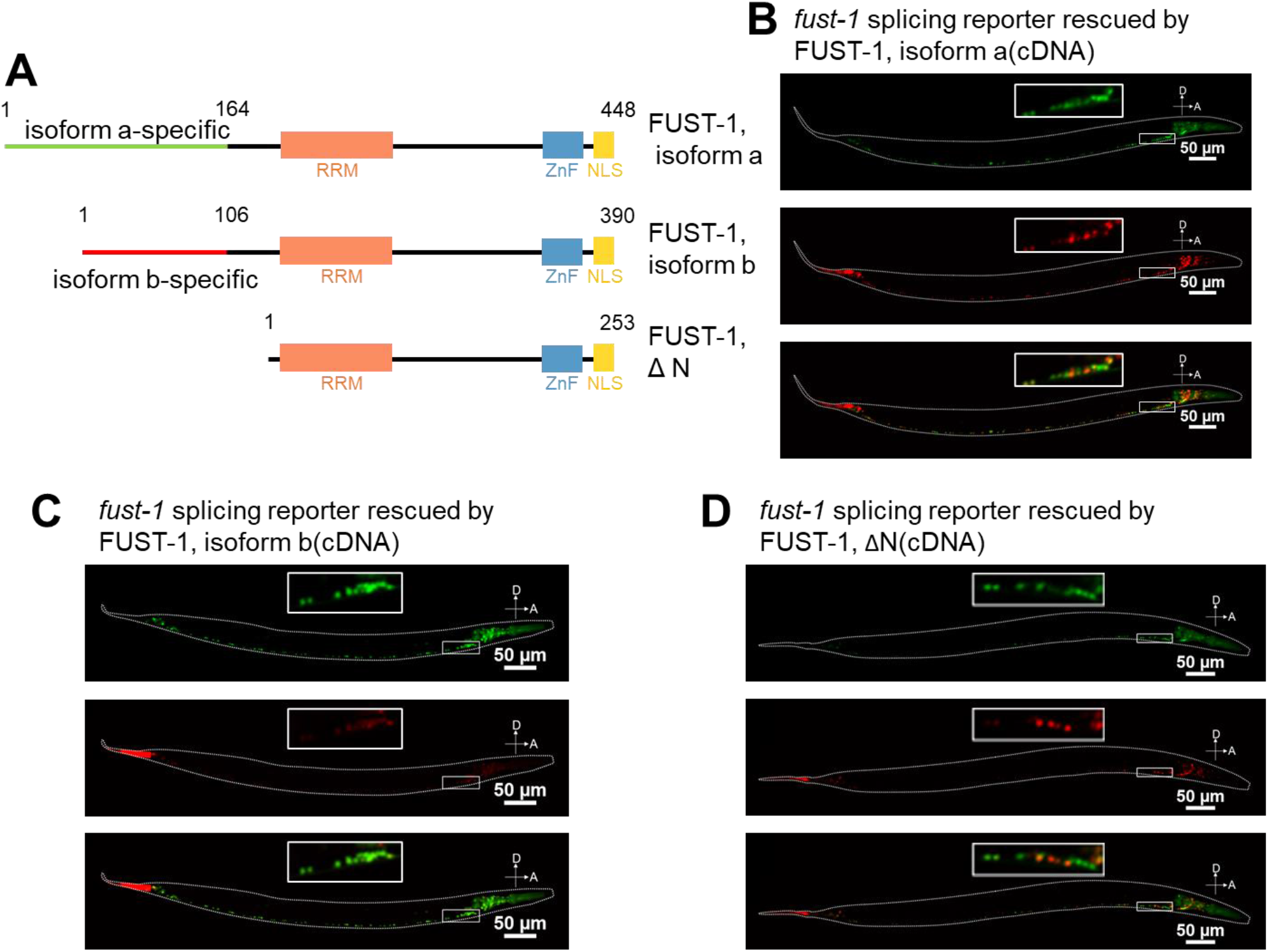
related to Figure 5. (A) Domains in three FUST-1 constructs. The lengths of the constructs are labeled. (B, C, and D) Representative confocal images showing rescue of *fust-1* exon 5 splicing reporter in FUST-1(csb21) strain by FUST-1, isoform a (B), FUST-1, isoform b (C), and by FUST-1, ΔN (D). Inset squares show the enlarged neck neurons in indicated strains. A: Anterior, D: Dorsal. Scale bars: 50 μm.

**Figure S6:**
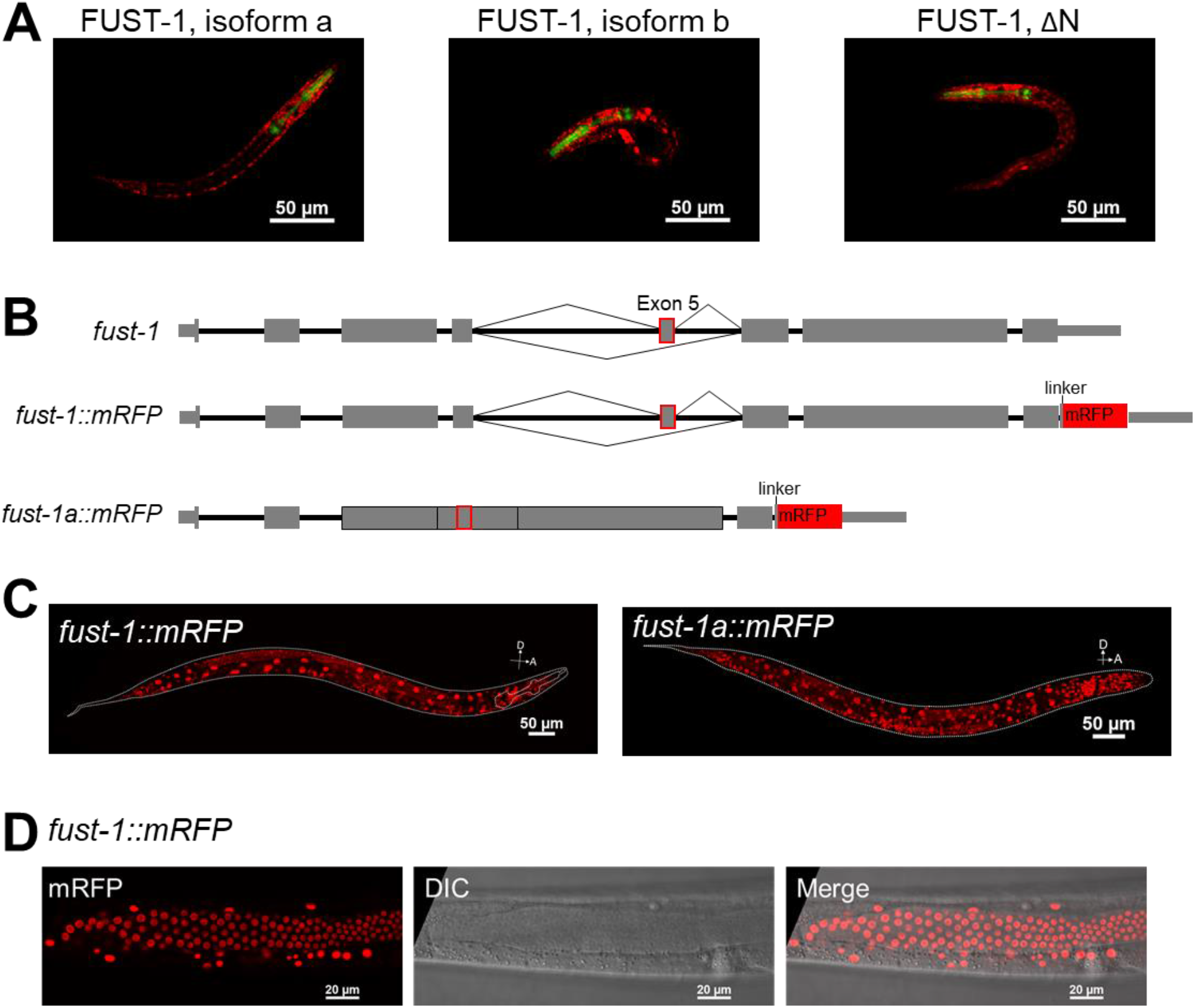
Related to Figure 6. (A) Representative confocal images of rescue strains with extrachromosomal expressions of mRFP fused cDNA of FUST-1 isoforms in the *fust-1(csb21)* strain. Note the expression of pharyngeal GFP. Worms were at the L1 stage. Scale bars: 50 μm. (B) Illustration of the gene structures of *fust-1* in wildtype N2 strain, C-terminal mRFP fused *fust-1 (fust-1::mRFP)* strain, and the strain can only express FUST1-, isoform a (*fust-1a::mRFP)*. (C) Representative images showing expression patterns of FUST-1 in the indicated strains. A: Anterior, D: Dorsal. Scale bars: 50 μm. (D) Confocal images showing that FUST-1 is expressed in the gonad. Scale bar: 20 μm.

